# Efficacy and gut dysbiosis of gentamicin-intercalated smectite as a new therapeutic agent against *Helicobacter pylori* in a mouse model

**DOI:** 10.1101/2020.06.30.179911

**Authors:** Su Jin Jeong, Kyoung Hwa Lee, Jie-Hyun Kim, Soon Young Park, Young Goo Song

## Abstract

**Background:** *Helicobacter pylori* eradication rate with conventional standard therapy is decreasing owing to antibiotic resistance, necessitating novel antibacterial strategies against *H. pylori*. We evaluated the efficacy of a gentamicin-intercalated smectite hybrid (S-GM)-based treatment, and analyzed fecal microbiome composition in *H. pylori*-infected mice.

**Methodology:** To evaluate anti-*H. pylori* efficacy, mice were divided into eight groups, and *H. pylori* eradication was assessed by *Campylobacter*-like organism (CLO) test and PCR assay of *H. pylori* in gastric mucosa. One week after *H. pylori* eradication, proinflammatory cytokine levels and atrophic changes in gastric mucosa were examined. Stool specimens were collected and analyzed for microbiome changes. The S-GM-based triple regimen decreased bacterial burden *in vivo*, compared with that in untreated mice or mice treated with other regimens. The therapeutic reactions in the CLO test from gastric mucosa were both 90% in standard triple therapy and S-GM therapy group, respectively. Those of *H. pylori* PCR in mouse gastric mucosa were significantly lower in standard triple therapy and S-GM therapy groups than in non-treatment group. Toxicity test results showed that S-GM therapy reduced IL-8 level and atrophic changes in gastric mucosa. Stool microbiome analysis revealed that compared with mice treated with the standard triple therapy, mice treated with the S-GM therapy showed microbiome diversity and abundant microorganisms at the phylum level.

**Conclusion:** Our results suggested that S-GM is a promising and effective therapeutic agent against *H. pylori* infection.

**Author summary:** The eradication rate on *Helicobacter pylori* (*H. pylori*) showed decreasing trend due to antibiotic resistance, especially clarithromycin. Therefore, we made a smectite hybrid as a drug delivery system using aminoglycosides antibiotic-gentamicin, and applied it to the mouse stomach wall to confirm the localized therapeutic effect, and set the different treatment duration to verify the effect. As a result, it was confirmed that the therapeutic efficacy of gentamicin (GM)-intercalated smectite hybrid (S-GM) was not inferior to the existing standard triple therapy, based on amoxicillin and clarithromycin, and preserved the diversity of gut microbiome composition. Therefore, a S-GM treatment is expected to be a new alternative regimen to *H. pylori* infection.

## Introduction

In 1983, Warren and Marchall described the gram-negative, spiral shaped microaerophilic bacterium *Helicobacter pylori* that colonizes the human stomach. *H. pylori* triggers numerous pathologic alterations in the stomach, including peptic ulcer disease, primary gastritis, and gastric cancer (1, 2). *H. pylori* eradication cures gastritis and alters the complication or recurrence of gastrointestinal diseases (3). The standard treatment for *H. pylori* infection is a triple therapy combining a proton pump inhibitor (PPI), clarithromycin, metronidazole, or amoxicillin (4). This regimen, however, fails to eradicate infection in 10–40% of patients and sometimes causes side effects (4-6). A major cause of this failure is the increase in multidrug-resistant *H. pylori* strains; hence, there is an alarming need to develop alternative antimicrobial agents with improved effectiveness.

Previously, we have confirmed that aminoglycosides have low minimum inhibitory concentration for recently isolated *H. pylori*, including major drug-resistant strains (7). However, aminoglycosides are polar, water-soluble compounds with very poor intestinal membrane permeability, resulting in low oral bioavailability (8, 9). Therefore, we used smectite clay, comprising tetrahedral sheets of SiO4 units and octahedral sheets of Al^3+^ ions (10), as a carrier of hydrophilic drugs to synthesize a gentamicin (GM)-intercalated smectite hybrid (S-GM) as a novel therapeutic agent. We previously identified that S-GM stably releases GM to the gastric wall, and a S-GM-based triple regimen decreases bacterial burden *in vivo* compared with that in untreated mice or mice treated with other regimens (11).

The human gut microbiota interacts with the host immune system and maintains metabolic homeostasis; thus, it is associated with obesity, inflammatory bowel disorder, allergic diseases, and neurological disorders (12, 13). Despite anatomical and compositional differences between human and mouse microbiota, some studies reported a concordance of microbiota shift in murine models and human diseases (14). Therefore, we analyzed changes in fecal microbiota in an *H. pylori*-infected murine model to examine the toxicity of S-GM. Because S-GM is not absorbed in the gastrointestinal tract, it is impossible to evaluate its pharmacokinetics (PK) and pharmacodynamics (PD). Moreover, in contrast to previous studies (11), S-GM was administered less frequently in this study.

Here, we aimed to evaluate the effect of dosing interval on daily administration of S-GM and to assess the safety of S-GM. Changes in inflammatory cytokine levels, atrophy of gastric mucosa, and fecal microbiota were analyzed after eradication of *H. pylori* with S-GM and compared with those after the standard triple therapy.

## Results

### CLO test and PCR assay of H. pylori in gastric mucosa

The S-GM-based regimen decreased *H. pylori* bacterial burden *in vivo*, compared with that in the untreated mice or mice treated with other regimens. Table 1 shows the therapeutic effect of each regimen on *H. pylori* infection. CLO test results showed that the therapeutic reactions in gastric mucosa were 90%, 90%, 80%, 80%, 70%, and 10% in Groups III, IV, V, VI, VII, and VIII, respectively (Table 1). The CLO scores of Groups III and IV were the lowest among the *H. pylori*-infected groups and were significantly lower than of Group II. The S-GM based therapy was not inferior to the standard triple therapy with amoxicillin and clarithromycin. Three or four doses per week also showed significant therapeutic results in the CLO test, although lower than that of daily administration.

**Table 1.** Individual data of CLO test of mouse gastric mucosa after treatment of HP infection.

| Group | Inoculation |  |  | Percentage of animals with positive<sup>a</sup><br>CLO test result, % |  |  | CLO scores |
| --- | --- | --- | --- | --- | --- | --- | --- |
|  | HP infection | Treatment | Duration of treatment | Positive | Partially positive | Negative |  |
| I | No | DW | D1-D7 | 0 | 0 | 100 | 0.0±0.0 |
| II | Yes | DW | D1-D7 | 80 | 20 | 0 | 2.8±0.4 |
| III | Yes | AMX + CLR + PPI | D1-D7 | 0 | 10 | 90 | 0.1±0.3* |
| IV | Yes | AMX + S-GM + PPI | D1-D7 | 0 | 10 | 90 | 0.1±0.3* |
| V | Yes | S-GM + PPI | D1-D7 | 0 | 20 | 80 | 0.2±0.4* |
| VI | Yes | AMX + S-GM + PPI | D1, D3, D5, D7 | 0 | 20 | 80 | 0.2±0.4* |
| VII | Yes | AMX + S-GM + PPI | D1, D4, D7 | 0 | 30 | 70 | 0.3±0.5* |
| VIII | Yes | AMX + S-GM + PPI | D1 | 60 | 30 | 10 | 2.3±1.1 |

HP, *Helicobacter pylori*; DW, distilled water; AMX, amoxicillin; CLR, clarithromycin; PPI, proton pump inhibitor; S-GM, gentamicin-intercalated smectite; CLO, *Campylobacter*-like organism
<sup>a</sup>A positive result indicates *H. pylori* colonization, which was observed as a color change in the medium from yellow to red.
\*Significantly different from Group II ( $P<0.01$ )

PCR assay was conducted to evaluate the therapeutic effects of S-GM in *H. pylori*-infected mice (Table 2). The amount of *H. pylori* DNA in mouse gastric mucosa was significantly lower in Groups III–VIII than in Group II.

**Table 2.**
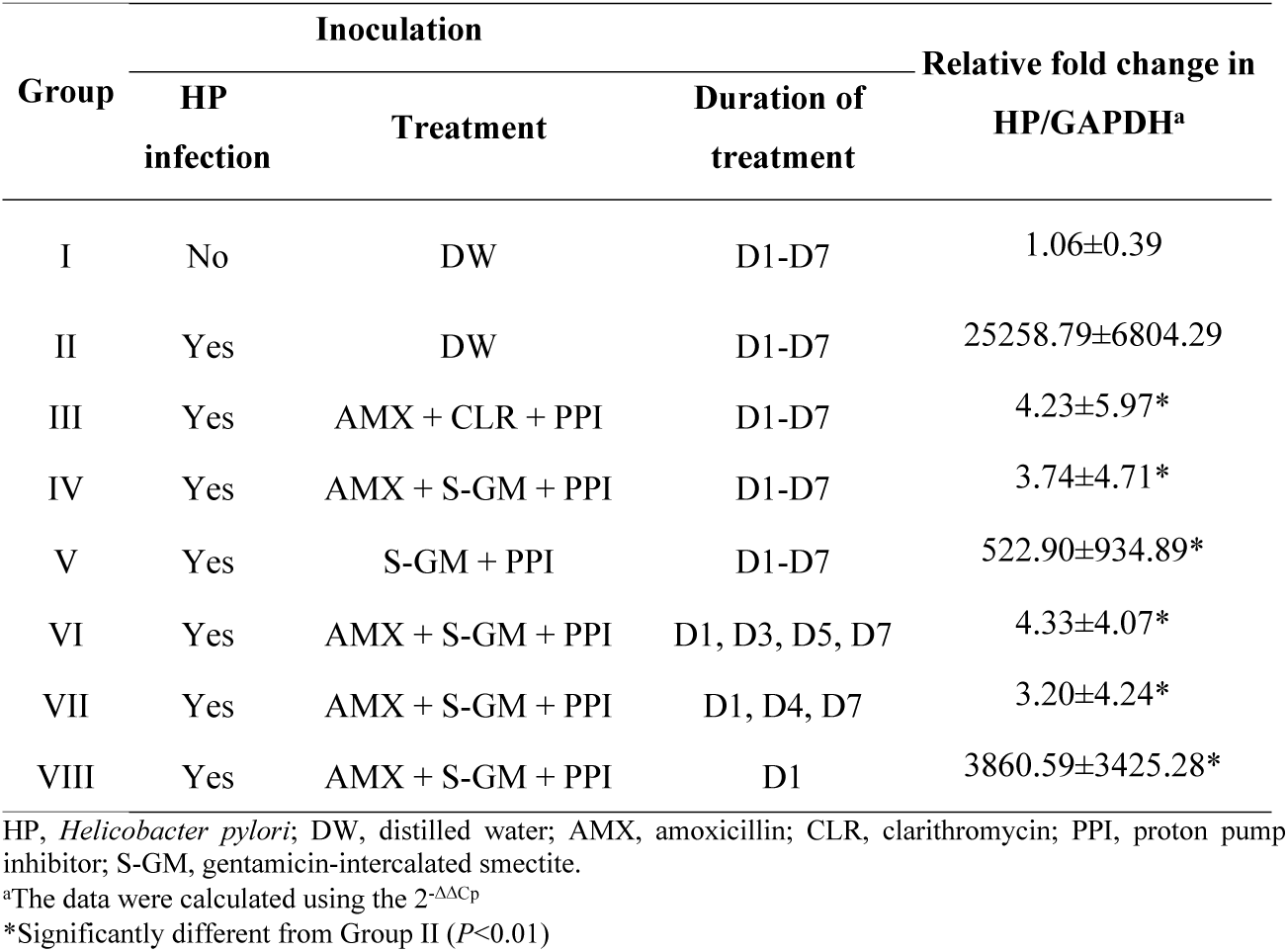
Individual data of quantitative PCR of mouse gastric mucosa after treatment of HP infection.

| Group | Inoculation |  |  | Relative fold change in HP/GAPDH<sup>a</sup> |
| --- | --- | --- | --- | --- |
|  | HP infection | Treatment | Duration of treatment |  |
| I | No | DW | D1-D7 | 1.06±0.39 |
| II | Yes | DW | D1-D7 | 25258.79±6804.29 |
| III | Yes | AMX + CLR + PPI | D1-D7 | 4.23±5.97* |
| IV | Yes | AMX + S-GM + PPI | D1-D7 | 3.74±4.71* |
| V | Yes | S-GM + PPI | D1-D7 | 522.90±934.89* |
| VI | Yes | AMX + S-GM + PPI | D1, D3, D5, D7 | 4.33±4.07* |
| VII | Yes | AMX + S-GM + PPI | D1, D4, D7 | 3.20±4.24* |
| VIII | Yes | AMX + S-GM + PPI | D1 | 3860.59±3425.28* |

HP, *Helicobacter pylori*; DW, distilled water; AMX, amoxicillin; CLR, clarithromycin; PPI, proton pump inhibitor; S-GM, gentamicin-intercalated smectite.
<sup>a</sup>The data were calculated using the $2^{-\Delta\Delta C_p}$
\*Significantly different from Group II ( $P<0.01$ )

### Proinflammatory cytokines and atrophy of gastric mucosa

S-GM-based therapy reduced IL-8 and TNF-α levels compared with the standard triple therapy (Group III). The degree of atrophic changes in gastric mucosa was analyzed in gastric tissue specimens; compared with the standard triple therapy (Group III), S-GM based therapy (Group IV) led to less atrophic changes in mouse stomach (Table 3).

**Table 3.** Plasma cytokine concentrations of IL-8 and TNF-α in each group.

| Group | HP Infection | IL-8 (pg/mL) | TNF- $\alpha$ (pg/mL) | Atrophy | Fecal microbiome | | | |
| --- | --- | --- | --- | --- | --- | --- | --- | --- |
|  |  |  |  |  | Alpha diversity |  | Abundance (%) |  |
|  |  |  |  |  | Shannon | Chao 1 | Bacteroidetes | Firmicutes |
| I | No | 17.13±5.66* | 225.00 ± 253.55 | 0.00±0.00 | 3.86±0.71 | 523.58±96.92 | 22.47 | 55.81 |
| II | Yes | 31.48±6.37 | 745.00 ± 485.64 | 1.60±0.52 | 3.78±0.60 | 483.81±81.67 | 28.84 | 53.87 |
| III | Yes | 18.60±9.06* | 564.50 ± 549.98 | 1.50±0.85 | 2.92±0.53** | 245.71±121.23** | 1.59** | 58.67 |
| IV | Yes | 14.70±6.70* | 442.50 ± 328.69 | 1.20±0.79 | 3.13±0.55 | 440.45±213.56† | 29.36† | 29.72 |

Data are expressed as mean ± standard error of 10 mice per group ( $\mu$ g/mL).
\*Significantly different from the positive control (Group II) ( $P$ <0.05)
\*\*Significantly different from Groups I and II ( $P$ <0.05)
†Significantly different from Group III ( $P$ <0.05)

### Changes in fecal microbiota

Formetagenomic analysis of S-GM, changes in the diversity and abundance of stool microbiome were identified (Table 3). Alpha diversity was the lowest in the standard triple therapy group (Group III). Shannon and Chao1 indices were relatively preserved in the S-GM therapy group (Group IV) compared with those in the standard triple therapy group (Group III vs. Group IV, Shannon index, 2.92 ± 0.53 vs. 3.13 ± 0.55; Chao 1, 245.71 ± 121.23 vs. 440.45 ± 213.56). Focusing on changes in abundance, the composition of stool microbiome was analyzed at the phylum and class levels. The abundance in Group III was significantly reduced, but preserved in Group IV with a similar trend as that in Group II (Figure 1 and 2).

**Figure 1.**
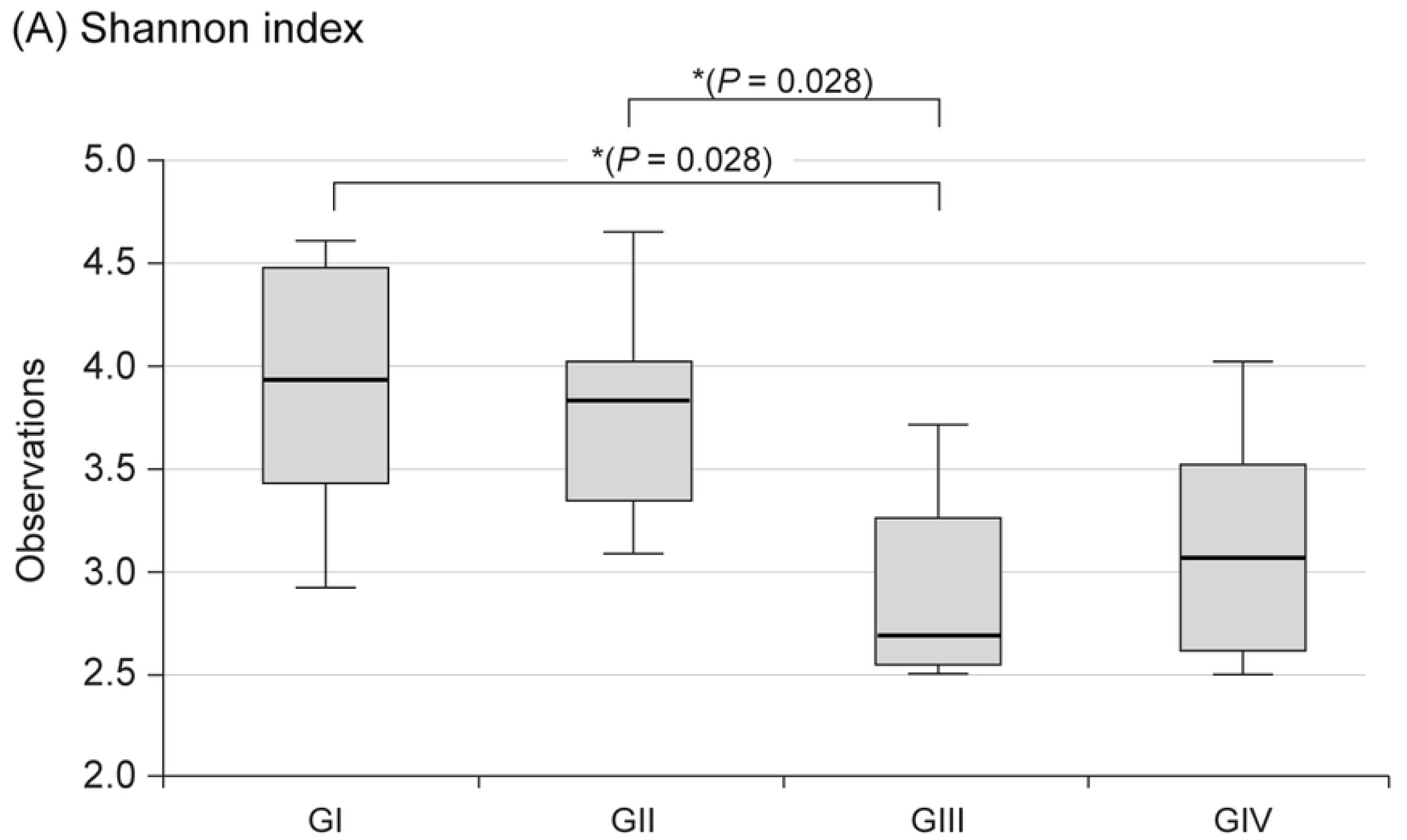
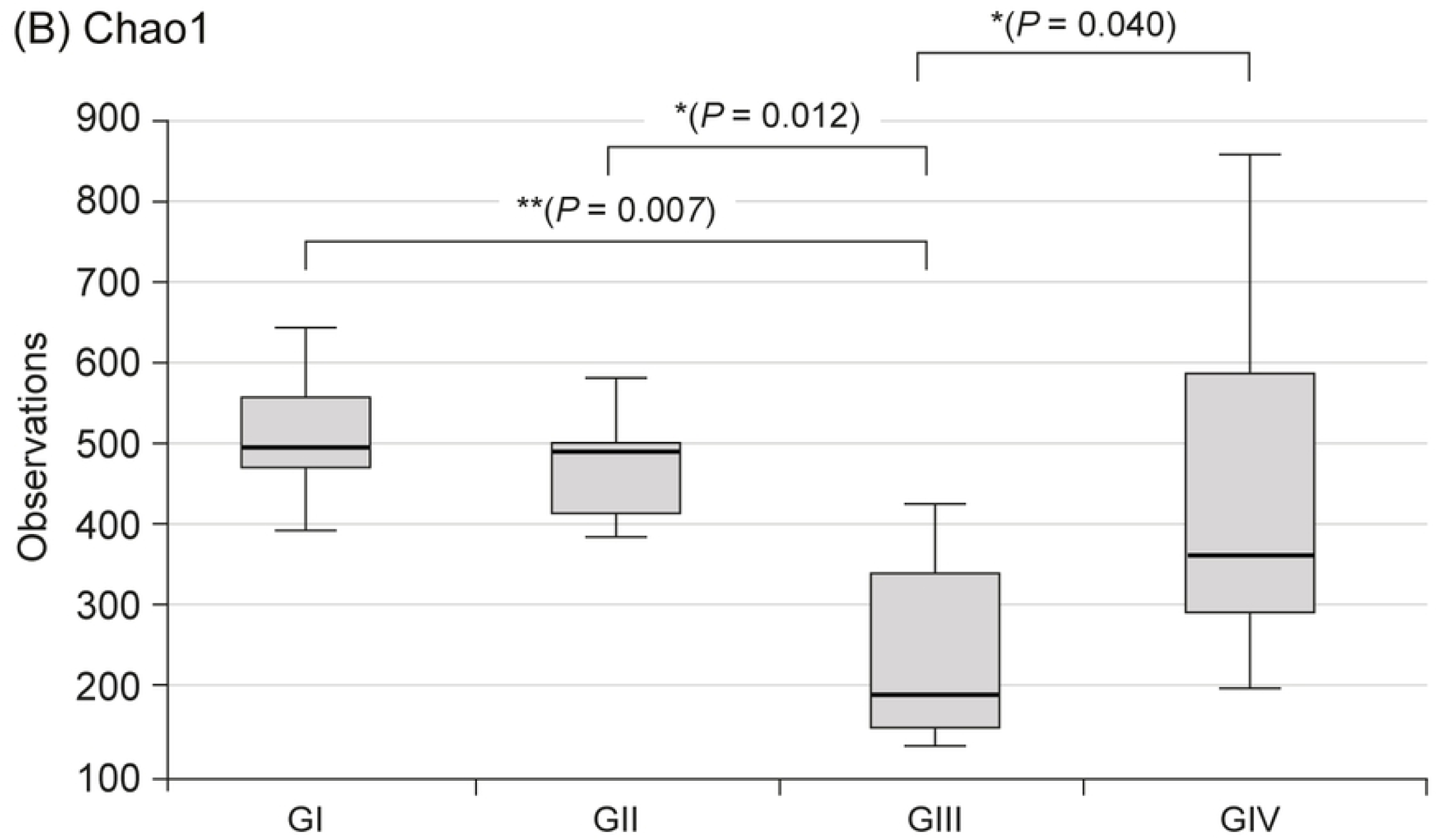
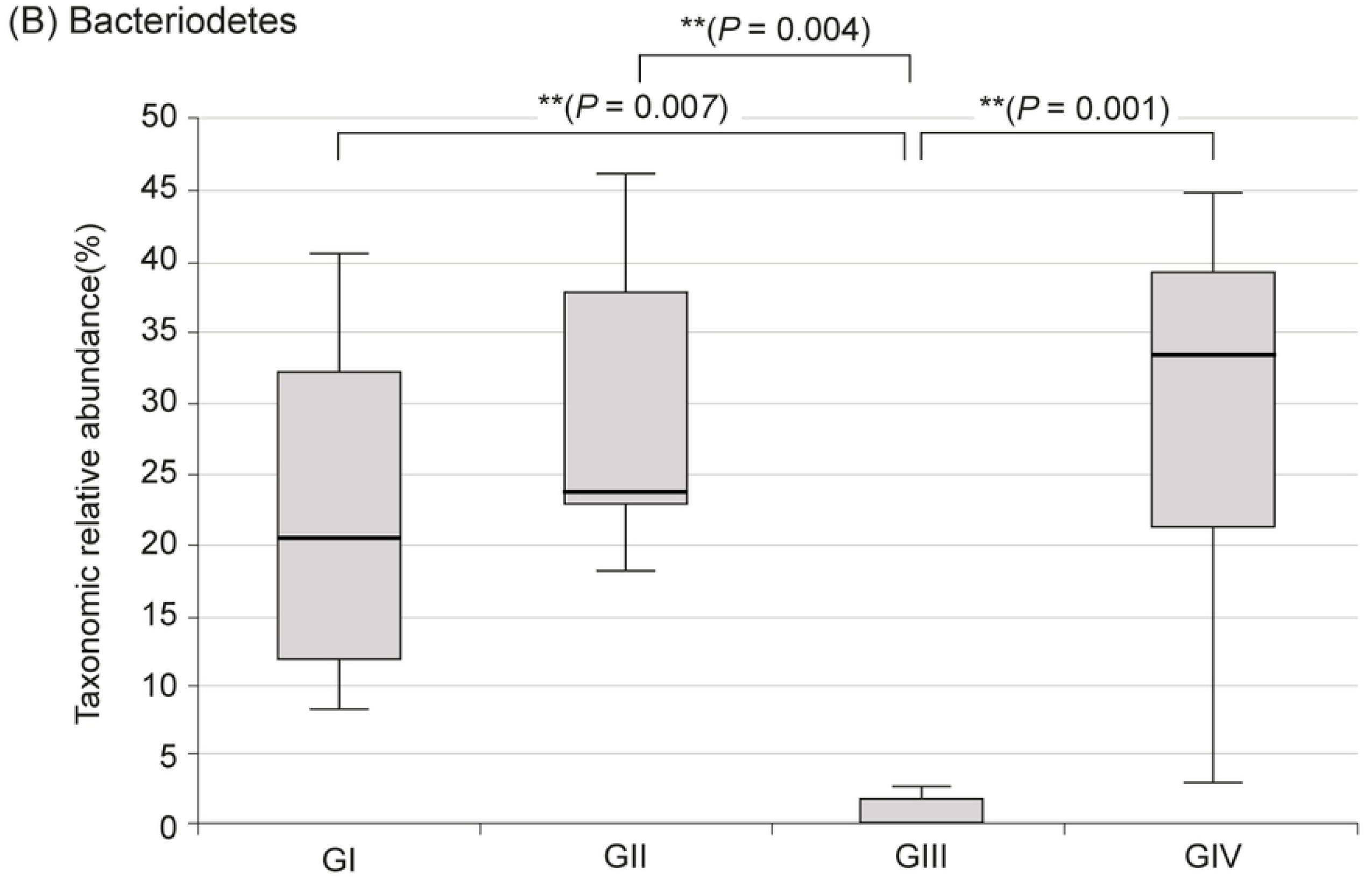
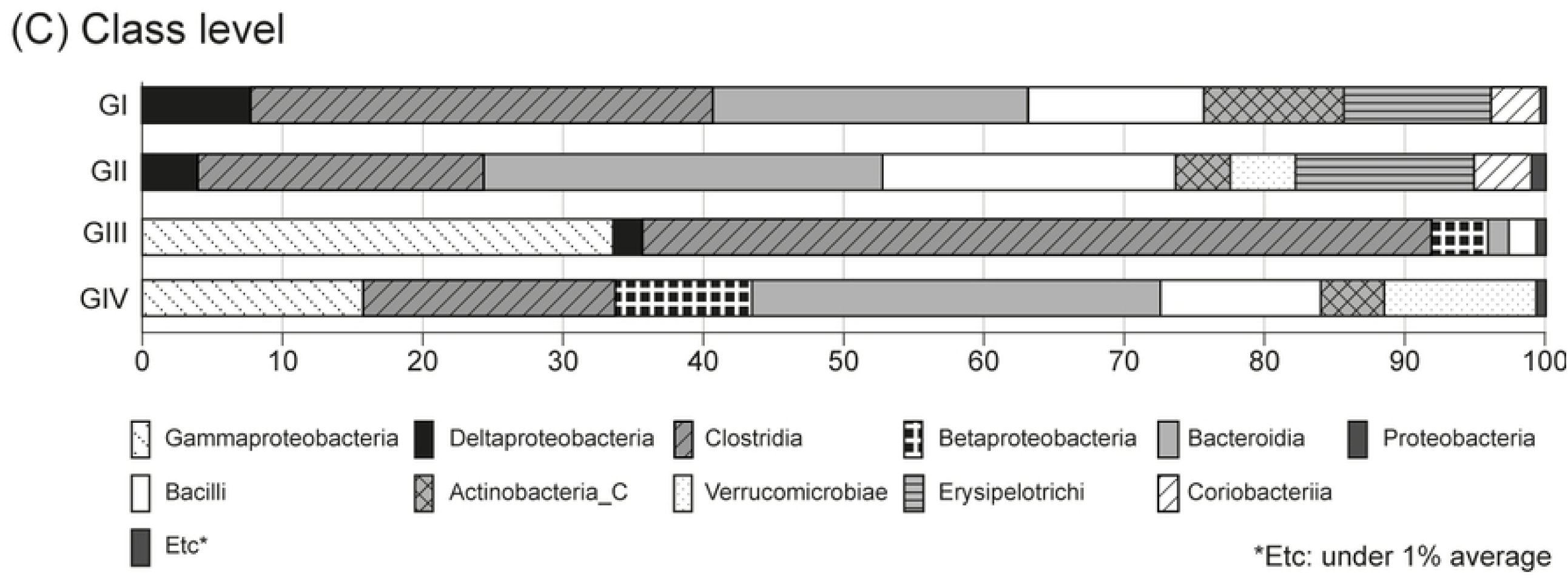
Alpha diversity in fecal microbiome between four groups.

**Figure 2.**
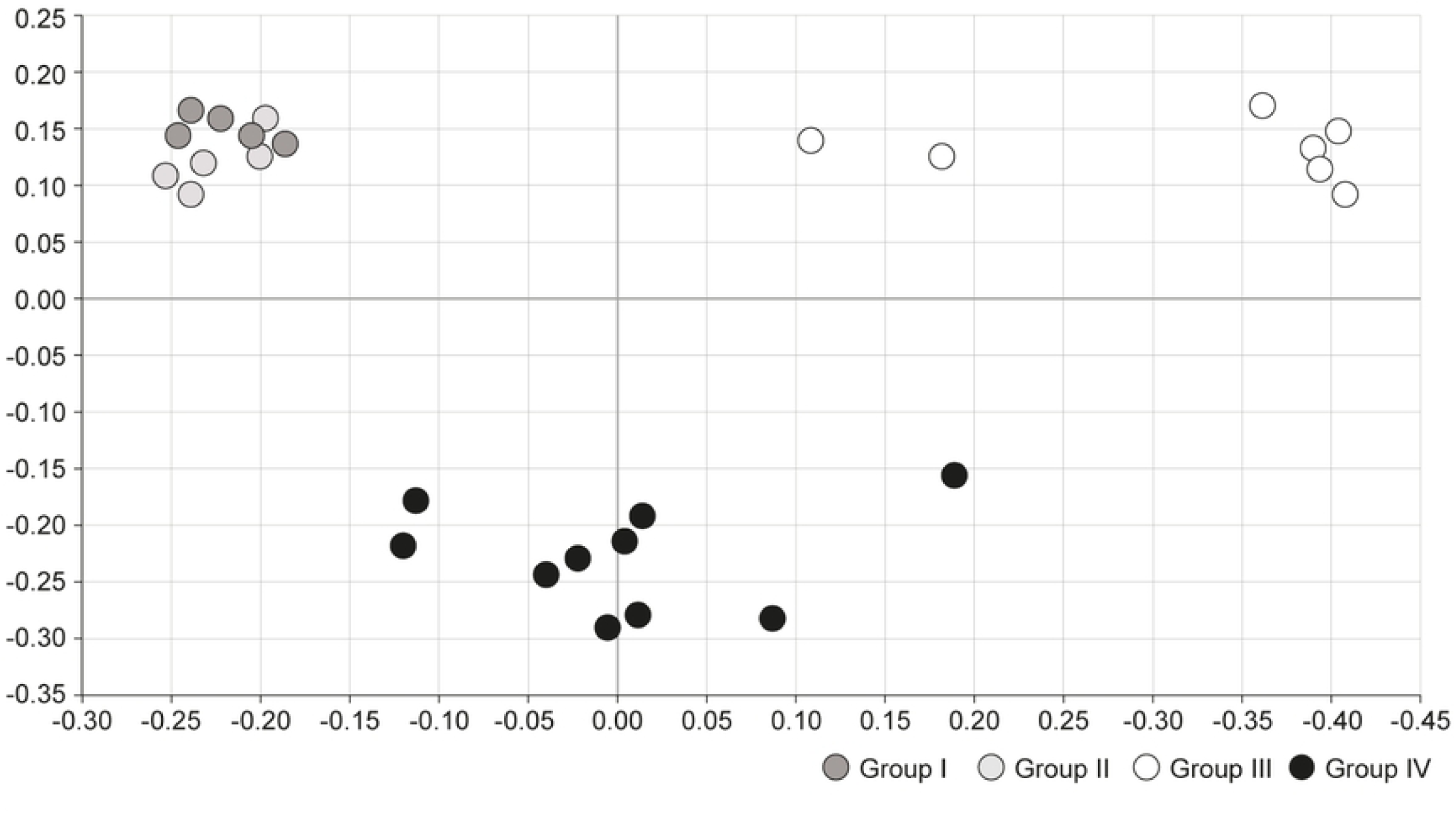
Microbiota composition and relative abundance distributions in four groups.

Principal coordinates analysis (PCoA) was conducted to compare microbial communities between the four groups. Group I and II showed similar trends, whereas Group III showed a distinctly different trend of microbiome composition. Group IV, which was treated with S-GM therapy, showed moderate disposition (Figure 3).

**Figure 3.**
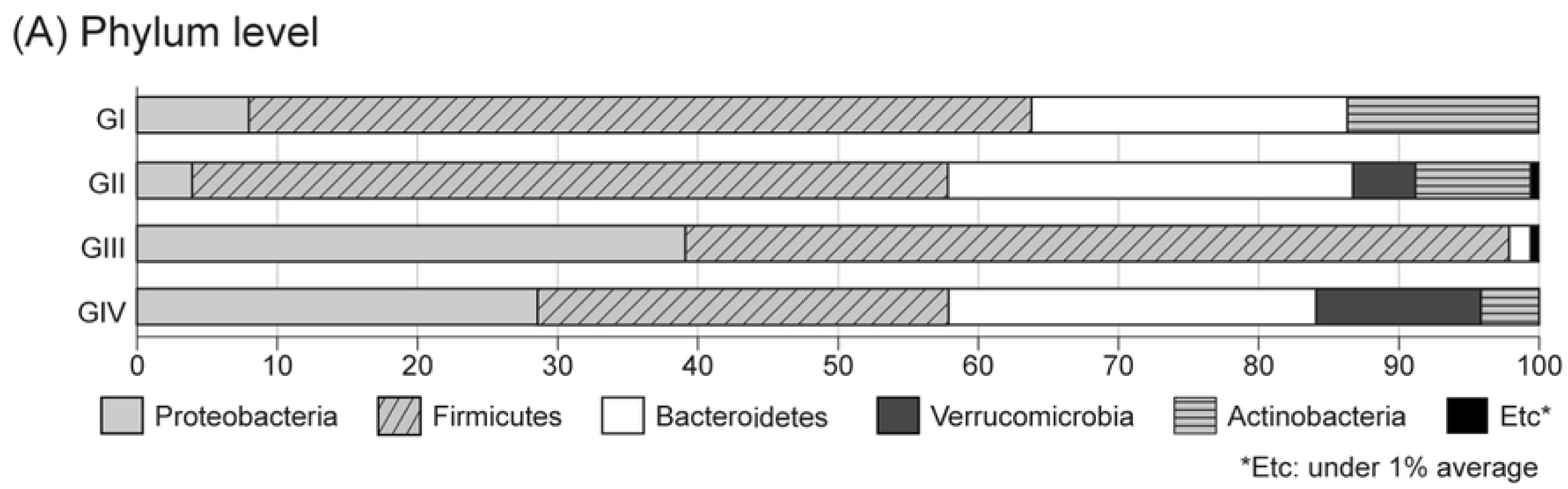
Comparison of microbial communities using principal coordinate analysis.

## Discussion

The existence of *H. pylori* in the human stomach has been known since as early as 60,000 years ago (15); it has been isolated from the gastric antrum and cultivated *in vitro* (16). Early eradication-based therapies regress *H. pylori*-associated diseases (3, 4). However, the eradication treatment efficacy has been compromised in many countries owing to the increasing resistance to antimicrobial agents (4-6, 17, 18). Additionally, recurrence of *H. pylori* remains a serious challenge worldwide, particularly in developing countries. The annual recurrence risk was 3.4% (95% CI, 3.1-3.7%) in high-income countries and 8.7% (95% CI, 8.8-9.6%) in low-income countries.

To improve the eradication efficacy, studies continue to evaluate novel treatment regimens, including quintuple therapies (19), high-dose dual therapies (20), and standard triple therapies with probiotics (21). However, the evidence is insufficient, and the side and cost effects of such therapies should be considered.

In our previous work, we demonstrated the high anti-*H. pylori* efficacy of S-GM in reducing *H. pylori* load in mouse stomachs (11). Here, we found no significant difference between daily administration and three- or four-time administration a week of S-GM. Therefore, we examined the possibility of administration three or four times a week for *H. pylori* eradication. However, the single dose therapy (Group VIII) showed significantly reduced therapeutic effect. Further, in future studies, S-GM efficacy should be confirmed with reduced overall treatment durations, such as 3, 5, and 7 days of daily treatment, not three or four times per week.

However, we could still assess the efficacy of S-GM in eradicating *H. pylori* in this study. GM concentration in the S-GM hybrid was intercalated only up to 8 mg/kg. If GM concentration is increased, or if it is intercalated with another drug delivery system similar to smectite capable of delivering antibiotics to the stomach wall, we can achieve prolonged and improved drug release. Intercalation of GM high concentrations is difficult to consider owing to the systemic side effects associated with intravenous administration, but its effectiveness can be expected in targeted localized therapy, such as *H. pylori* eradication. Further research is needed to compare the therapeutic effect of S-GM with other drug delivery systems, such as alginate and a composite. With chitosan-treated beads, alginated-antibiotics hybrids may achieve pH-dependent retarded release of highly soluble drug (22).

In the present study, S-GM triple therapy reduced IL-8 level and atrophic changes in gastric mucosa. Further, stool microbiome analysis showed that microbiome diversity and microorganism abundance at the phylum level were preserved in the S-GM triple therapy group. PK/PD analysis is required to examine the toxicity of S-GM; however, because S-GM is not absorbed systemically, PK/PD analysis is not feasible for S-GM. S-GM as a localized therapy showed bactericidal effect against *H. pylori* attached to the gastric wall. Therefore, toxicity analysis of this treatment was focused on changes in intestinal bacterial microbiome, and the results confirmed that the components of microbiome were well preserved compared with those after the standard therapy.

In this study, amoxicillin- and clarithromycin-based standard therapies have been shown to lead to microbiome dysbiosis, which is associated with various metabolic diseases, gastrointestinal diseases, and even gastric cancer. *H. pylori* infection itself as well as decreased microbial diversity and abundance are correlated with gastric carcinoma (23, 24). Therefore, our results indicated that S-GM therapy will not only block gastric carcinogenesis but also reduce the incidence of diseases, such as inflammatory bowel and metabolic diseases, by minimalizing changes in gut microbiota with low toxicity in addition to sufficient efficacy compared with standard therapy.

A previous *H. pylori* and microbiome study revealed a dramatic decrease in microbiome diversity immediately within 1 week after eradication, indicating that the bacterial community resembled and recovered the pre-antibiotic period only 4 years after a long-term follow-up. In humans, *Actinobacteria* was the most affected by antibiotics (25). In our mouse model study, *Actinobacteria* reduction was also noticeable after the use of amoxicillin and clarithromycin (Group III). Thus, to reduce prolonged dysbiosis and its consequences, it is necessary to eradicate *H. pylori* with minimal use of antibiotics, for which S-GM may be an effective strategy.

Therefore, to decrease antibiotic-related gut dysbiosis in patients and maintain microbiome components, targeted therapy for *H. pylori* attached to the gastric wall is needed instead of therapy with systemic antibiotics. Moreover, the use of a smectite applied to the stomach wall as a drug delivery system would be a significant turning point for *H. pylori* eradication.

There were, however, several limitations in this study. First, the study used animal models; thus, the actual clinical dysbiosis may differ in humans. Second, the stool microbiome analysis was not conducted for each individual mouse, and stool was extracted within the same treatment group. The mouse itself could share the same microbiome environment owing to co-housing in the same cage (26, 27). Therefore, for more accurate analysis, it is necessary to analyze feces of each mouse subject and compare the individual eradication rate with a specific therapeutic regimen. Third, long-term follow-up after S-GM treatment is needed.

Nevertheless, this is the first study to verify the gut dysbiosis of the S-GM as an alternative therapy for *H. pylori* eradication to overcome the increasing antibiotic resistance to other regimens. Moreover, localized *H. pylori* eradication will make a novel paradigm shift in *H. pylori* treatment.

## Materials and Methods

### Intercalation of GM

GM (2 mg/mL) solution was prepared using gentamicin sulfate of USP grade produced by BIO BASIC INC (Toronto, Canada). Ca-smectite was prepared by purifying the bentonite found in the area of Gampo, Korea. To generate a GM-intercalated smectite hybrid, GM solution was mixed with Ca-smectite to a concentration of 250 mL/g, and the mixture was stirred vigorously for 24 h. Next, the hybrid solution was dialyzed with 5 L of distilled water (DW) for ∼8 h at 50°C, and the dialysis was repeated three to four times until sulfate ions could not be detected by PbCl2. A hybrid powder was finally obtained by frieze-drying the dialyzed hybrid solution for 2–3 days. The amount of GM released from the hybrid was determined by batch-release test using 25 mL of pH 1.2 solution for 100 mg of the hybrid powder. The total amount of GM released within 1 h was determined to be ∼5.0 mg per 100 mg of the hybrid.

### Animal preparation

The Institutional Animal Care and Use Committee at Daegu-Gyeongbuk Medical Innovation Foundation (DGMIF), Daegu, Korea, approved the animal procedures. Four-week-old male C57BL/6 mice were purchased from Japan SLC, Inc., Shizuoka, Japan. The mice were 5 weeks of age and weighed 18–20 g at the start of the experiment. The animal experiments were reviewed and approved by the Institutional Animal Care and Use Committee of the DGMIF.

### Anti-H. pylori efficacy in vivo

#### H. pylori strains and culture conditions

*H. pylori* SS1 was used in this study. The bacteria were maintained and grown on Brucella agar (Merck, Germany) supplemented with 10% fetal bovine serum (Gibco, USA), and incubated under microaerobic conditions (5% O2, 10% CO2, and 85% N2) at 37 °C for 72 h.

#### Inoculation of experimental animals

For *in vivo* assessment of anti*-H. pylori* effect, 80 mice were allowed to acclimatize for 1 week before the initiation of the experiment. After the acclimatization period, the animals were fasted for 12 h, and 70 of them were intragastrically infected with 0.5 mL of 2.0×10^9^ CFU/mL *H. pylori* suspension by oral gavage every 48 h, and this was repeated three times in 1 week.

#### Distribution of animals

A total of 80 mice were used for analysis, and 70 *H. pylori-*infected mice were distributed into seven groups and allowed to rest for 1 week after the last inoculation. Group I was a normal group consisting of uninfected mice. Group II, a negative control group, received DW as a vehicle. Group III, a positive control group, was treated with the standard triple therapy consisting of amoxicillin (AMX) (14.25 mg/kg), clarithromycin (CLR) (14.3 mg/kg), and a PPI (omeprazole 138 mg/kg). Group IV was treated with AMX (14.25 mg/kg), S-GM (which emitted 8 mg/kg of GM), and a PPI (138 mg/kg). Group V was treated with S-GM (which emitted 8 mg/kg of GM) and a PPI (138 mg/kg). Groups V–VIII were treated with the same regimen as that of Group IV, but with different administration intervals of four times per week, three times per week, and a single dose per week, respectively. In Groups I–IV, the treatments were orally administered to mice once a day for 7 consecutive days. The *H. pylori*-IgG level was checked with an ELISA kit (Cusabio Biotech Co., USA) before the treatment period to confirm the serological status of *H. pylori*-infected mice.

#### CLO test and PCR assay of H. pylori in gastric mucosa

At 12 h after the last administration, mice were euthanized, and their stomachs were removed from their abdominal cavities. Samples of gastric mucosa from the pyloric region were assayed with CLO kits (Asan Pharmaceutical Co., Seoul, Korea) and incubated at 37°C for 12 h to examine urease activity. The reaction score was graded from 0 to 3 with 0 = no color change, 1 = bright red, 2 = light purple, and 3 = dark red.

*H. pylori* DNA was prepared using the bead beater-phenol extraction method(28). A bacterial suspension was placed in a 2.0-mL screw-cap microcentrifuge tube filled with glass beads (Biospec Products, Bartlesville, OK, USA) and 200 μL of phenol:chloroform:isoamyl alcohol solution (50:49:1). After an initial denaturation/activation step (95°C for 5 min), DNA (50 ng) was amplified in a 20-μL volume for 35 cycles of denaturation (94°C for 60 s), annealing (62°C for 60 s), and extension (72°C for 90 s) using the following primers: *H. pylori*-specific *ureA* and *ureC*, sense, 5′-TGATGCTCCACTACGCTGGA-3′, and antisense, 5′-GGGTATGCACGGTTACGAGT-3′ (expected product 265 bp); (29) and GAPDH, sense, 5′-TGGGGTGATGCTGGTGCTG-AG-3′, and antisense, 5′-GGTTTCTCCAGGCGGCATGTC-3′ (expected product 497 bp)(30). The PCR products were analyzed by electrophoresis in 1.5% agarose gels.

#### Proinflammatory cytokines and atrophy of gastric mucosa

Plasma was obtained on day 21 through insertion of a heparinized microhematocrit tube into the ophthalmic venous plexus of mice. Plasma IL-8 and TNF-α levels were measured using mouse ELISA kits (R&D System, Minneapolis, MN, USA).

For histopathologic analysis, the stomach was fixed in 10% neutralized buffered formalin, and embedded in paraffin. Sections (4-μm thick) were then stained with hematoxylin and eosin. The glandular mucosae of the corpus and antrum were examined histologically. Atrophic changes, as defined by atrophy of glandular cells and hyperplasia of mucus cells, were determined in a blinded fashion and scored based on the percentage of altered gastric mucosa(31): 0 = no mucosal alterations; 1 = less than 5%; 2 = 10% to 25%; 3 = 25% to 50%; and 4 = 50% to 75%.

### Fecal microbiota

#### DNA extraction from fecal materials

After examination of IgG level post-treatment, the mice were sacrificed. Their feces were collected and frozen at −80°C until processed. From the fecal materials of each mouse, DNA was extracted using FastDNA^®^ SPIN Kit (MP Biomedicals, Solon, OH, USA). The samples were lysed with FastPrep^®^ Instruments and centrifuged, and DNA was isolated from the supernatant using the procedure of silica-based GENECLEAN^®^ and SPIN filters (MP Biomedicals, Solon, OH, USA)(32).

#### PCR amplification and 16S rRNA gene sequencing

Using the extracted metagenomics DNA as a template, PCR was performed for amplification of the V3-V4 regions of the bacterial 16S rRNA gene using the primers 341F (5′-TCGTCGGCAGCGTC-AGATGTGTATAAGAGACAG-CCTACGGGNGGCWGCAG-3′; the underlined sequence indicates the target region primer) and 805R (5′-GTCTCGTGGGCTCGG-AGATGTGTATAAGAGACAG-ACTACHVGGGTATCTAATCC-3′). Next, secondary amplification for attachment of the Illumina NexTera barcode was performed using the following primers (X indicates the barcode region): i5 forward primer, 5′-AATGATACGGCGACCACCGAGATCTACAC-XXXXXXXX-TCGTCGGCAGCGTC-3′; and i7 reverse primer, 5′-CAAGCAGAAGACGGCATACGAGAT-XXXXXXXX-AGTCTCGTGGGCTCGG-3′. The PCR products were identified via 1% agarose gel electrophoresis and visualized in a Gel Doc system (BioRad, Hercules, CA, USA).

After purification of the amplified products using Clean PCR (CleanNA, Waddinxveen, Netherlands), qualified products were assessed on a Bioanalyzer 2100 (Agilent, Palo Alto, CA, USA). The libraries were prepared for analysis, and gene sequencing was performed using an Illumina MiSeq Sequencing system (Illumina, San Diego, CA, USA) according to the manufacturer’s instructions.

#### Bioinformatics for microbiota analysis

The EzBioCloud 16S rRNA database (https://www.ezbiocloud.net) operated by ChunLab (ChunLab, Inc., Seoul, Korea) was used as a bioinformatics cloud platform for accurate pairwise and taxonomic assignments. Chimeric reads were filtered on reads with <97% similarity based on the UCHIME algorithm (33), and operational taxonomic units (OTU)s with single and un-clustered reads are omitted from further analysis. Alpha diversity, which measures the diversity and abundance of bacterial species, was analyzed by the Shannon and Chao1 indices (34). Beta diversity was measured using PCoA derived from Jensen-Shannon (35). The Wilcoxon rank-sum test was used to examine differences in the number of OTUs.

### Statistical analysis

Data are presented as means ± standard error, and the non-parametric Mann–Whitney test was used to compare groups. Multiple differences between groups were evaluated using one-way ANOVA multiple comparison test. The 95% confidential interval (CI) of the detection rate was obtained using the MINITAB statistical software (Minitab, Inc., State College, PA, USA). If two values were not overlapped between its 95% CI, the difference was considered significant. A *p* value of < 0.05 was considered statistically significant. Results were analyzed using the Statistics Package for Social Science (SPSS 15.0 for Windows; SPSS Inc., Chicago, IL, USA).

## Acknowledgement

We thank Myoung Ju Choi, Kwang Hoon Lee, and Jae Min Kim for helpful research support and discussions.

## Author contributions

SJJ, KHL, JK and YGS conceived and designed the research, SYP performed the experiments. JHK, and YGS analyzed the data, SJJ and KHL wrote the draft, and YGS were involved in the review and editing of the final manuscript. All authors read and approved the final manuscript.

## Funding

This work was supported by the Basic Research Project (Study No. GP2017-020) of the Korea Institute of Geoscience and Mineral Resources (KIGAM), funded by the Ministry of Science, ICT, and Future Planning of Korea. This work was also supported by the Seyoung Association (SYH) research grant from the Department of Internal Medicine, Gangnam Severance Hospital (2017-GNS-001).

## Competing interests

All authors report no potential conflicts of interest.

## Reference

1. L. M. Coussens, Z. Werb, Inflammation and cancer. Nature 420, 860–867 (2002).

2. J. G. Kusters, A. H. van Vliet, E. J. Kuipers, Pathogenesis of Helicobacter pylori infection. Clin. Microbiol. Rev. 19, 449–490 (2006).

3. P. Malfertheiner et al., Management of Helicobacter pylori infection-the Maastricht V/Florence Consensus Report. Gut 66, 6–30 (2017).

4. N. R. O’Morain, M. P. Dore, A. J. O’Connor, J. P. Gisbert, C. A. O’Morain, Treatment of H elicobacter pylori infection in 2018. Helicobacter 23, e12519 (2018).

5. N. O. Kaakoush, C. Asencio, F. Mégraud, G. L. Mendz, A redox basis for metronidazole resistance in Helicobacter pylori. Antimicrob Agents Chemother (Bethesda) 53, 1884–1891 (2009).

6. Z. Song et al., Prospective multi-region study on primary antibiotic resistance of Helicobacter pylori strains isolated from Chinese patients. Dig. Liver Dis. 46, 1077–1081 (2014).

7. K. H. Lee et al., Can aminoglycosides be used as a new treatment for Helicobacter pylori? In vitro activity of recently isolated Helicobacter pylori. Infect. Chemother. 51 (2019).

8. C. E. Cox, Gentamicin. Med. Clin. North Am. 54, 1305–1315 (1970).

9. J. Recchia, M. H. Lurantos, J. A. Amsden, J. Storey, C. R. Kensil, A semisynthetic Quillaja saponin as a drug delivery agent for aminoglycoside antibiotics. Pharm. Res. 12, 1917–1923 (1995).

10. F. Thomas et al., Layer charge and electrophoretic mobility of smectites. Colloids and Surfaces A: Physicochemical and Engineering Aspects 159, 351–358 (1999).

11. S. J. Jeong et al., Gentamicin-intercalated smectite as a new therapeutic option for Helicobacter pylori eradication. Journal of antimicrobial chemotherapy 73, 1324–1329 (2018).

12. J. K. Nicholson et al., Host-gut microbiota metabolic interactions. Science 336, 1262–1267 (2012).

13. C. A. Lozupone, J. I. Stombaugh, J. I. Gordon, J. K. Jansson, R. Knight, Diversity, stability and resilience of the human gut microbiota. Nature 489, 220–230 (2012).

14. T. L. Nguyen, S. Vieira Silva, A. Liston, J. Raes, How informative is the mouse for human gut microbiota research? Dis. Model. Mech. 8, 1–16 (2015).

15. Y. Moodley et al., Age of the association between Helicobacter pylori and man. PLoS Pathog. 8, e1002693–e1002693 (2012).

16. J. R. Warren, B. Marshall, Unidentified curved bacilli on gastric epithelium in active chronic gastritis. Lancet (British edition) 1, 1273–1275 (1983).

17. Miftahussurur et al., Surveillance of Helicobacter pylori Antibiotic Susceptibility in Indonesia: Different Resistance Types among Regions and with Novel Genetic Mutations. PLoS ONE 11, e0166199–e0166199 (2016).

18. E. N. Ontsira Ngoyi et al., Molecular Detection of Helicobacter pylori and its Antimicrobial Resistance in Brazzaville, Congo. Helicobacter 20, 316–320 (2015).

19. N. de Bortoli et al., Helicobacter pylori eradication: a randomized prospective study of triple therapy versus triple therapy plus lactoferrin and probiotics. Am. J. Gastroenterol. 102, 951–956 (2007).

20. J. Yang et al., High-dose dual therapy is superior to standard first-line or rescue therapy for Helicobacter pylori infection. Clin. Gastroenterol. Hepatol. 13, 895-905.e895 (2015).

21. B. Oh et al., The Effect of Probiotics on Gut Microbiota during the Helicobacter pylori Eradication: Randomized Controlled Trial. Helicobacter 21, 165–174 (2016).

22. H. H. Tønnesen, J. Karlsen, Alginate in drug delivery systems. Drug Dev. Ind. Pharm. 28, 621–630 (2002).

23. R. M. Ferreira et al., Gastric microbial community profiling reveals a dysbiotic cancer-associated microbiota. Gut 67, 226–236 (2018).

24. H. J. Jo et al., Analysis of Gastric Microbiota by Pyrosequencing: Minor Role of Bacteria Other Than Helicobacter pylori in the Gastric Carcinogenesis. Helicobacter 21, 364–374 (2016).

25. H. E. Jakobsson et al., Short-term antibiotic treatment has differing long-term impacts on the human throat and gut microbiome. PLoS ONE 5, e9836–e9836 (2010).

26. Z. Shi et al., Segmented Filamentous Bacteria Prevent and Cure Rotavirus Infection. Cell 179, 644-658.e613 (2019).

27. S. J. Robertson et al., Comparison of Co-housing and Littermate Methods for Microbiota Standardization in Mouse Models. Cell Reports 27, 1910-1919.e1912 (2019).

28. B.-J. Kim et al., Identification of mycobacterial species by comparative sequence analysis of the RNA polymerase gene (rpoB). J. Clin. Microbiol. 37, 1714–1720 (1999).

29. Y. B. Kim et al., The influence of number of gastroscopic biopsy specimens on follow-up Campylobacter-Like Organism (CLO) test. Korean J. Gastroenterol. 35, 422–428 (2000).

30. P. Kundu et al., Cag pathogenicity island-independent up-regulation of matrix metalloproteinases-9 and −2 secretion and expression in mice by Helicobacter pylori infection. Journal of biological chemistry 281, 34651–34662 (2006).

31. J. G. Fox et al., Concurrent enteric helminth infection modulates inflammation and gastric immune responses and reduces helicobacter-induced gastric atrophy. Nat. Med. 6, 536–542 (2000).

32. A. Layton et al., Development of Bacteroides 16S rRNA gene TaqMan-based real-time PCR assays for estimation of total, human, and bovine fecal pollution in water. Appl. Environ. Microbiol. 72, 4214–4224 (2006).

33. R. C. Edgar, B. J. Haas, J. C. Clemente, C. Quince, R. Knight, UCHIME improves sensitivity and speed of chimera detection. Bioinformatics 27, 2194–2200 (2011).

34. A. D. Willis, Rarefaction, alpha diversity, and statistics. Front. Microbiol. 10, 2407 (2019).

35. J. Lin, Divergence measures based on the Shannon entropy. IEEE Trans. Inf. Theory 37, 145–151 (1991).

